# Electrophysiological correlates of (mis)judging social information

**DOI:** 10.1101/2023.06.02.543470

**Authors:** Miles Wischnewski, Michael O.Y. Hörberg, Dennis J.L.G. Schutter

**Author notes:** Corresponding author: Miles Wischnewski University of Minnesota, Department of Biomedical Engineering Nils Hasselmo Hall, room 7-105, 312 Church St SE, Minneapolis, MN 55455.

## Abstract

Social information can be used to optimize decision making. However, the simultaneous presentation of multiple sources of advice can lead to a distinction bias in judging the validity of the information. While involvement of event-related potential (ERP) components in social information processing has been studied, how they are modulated by (mis)judging advisor’s information validity remains unknown. In two experiments participants performed a decision making task with highly accurate or inaccurate cues. Each experiment consisted of a initial, learning and test phase. During the learning phase three advice cues were simultaneously presented and the validity of them had to be assessed. The effect of different cue constellations on ERPs was investigated. In the subsequent test phase, the willingness to follow or oppose an advice cue was tested. Results demonstrated the distinction bias with participants over or underestimating the accuracy of the most uncertain cues. The P2 amplitude was significantly increased during cue presentation when advisors were in disagreement as compared to when all were in agreement, regardless of cue validity. Further, a larger P3 amplitude during outcome presentation was found when advisors were in disagreement and increased with more informative cues. As such, most uncertain cues were related to the smallest P3 amplitude. Findings suggest that misjudgment of social information is related to P3 amplitude subserving evaluation information and learning. This study provides novel insights into the role of P2 and P3 components during judgement of social information validity.

## Introduction

During uncertain decision-making individuals often seek guidance from external social information (Adolphs, 2009; Behrens et al., 2008; De Jaegher et al., 2010; Frith & Frith, 2012; Pescetelli et al., 2021; Suzuki et al., 2015). One example of such external information is the advice from individuals who we consider to be more knowledgeable than ourselves (Behrens et al., 2008; Diaconescu et al., 2017; Meshi et al., 2012; Wischnewski et al., 2018; Wischnewski & Schutter, 2019). When judging the usefulness of social information the dissonance between advisor opinions and oneself or others is assessed (Diaconescu et al., 2017; Meshi et al., 2012). Advice may seem more valid if it conquers with one’s own opinion or the opinion of other advisors that are deemed accurate. The situation becomes more complicated when conflicting social information from different sources is presented (Collins et al., 2011; De Martino et al., 2017; Hsee & Zhang, 2004). Complex strategies are used that consider one’s own uncertainty and estimate validity of different advisors, which in conjunction allows us to optimize our internal prediction model (Burnside & Ullsperger, 2020; De Martino et al., 2017; De Martino & Cortese, 2023).

Internal prediction models are a key aspect of performance monitoring, which refers to the a set of cognitive processes involved in tracking accuracy of decisions and updating behavior when errors are noticed (Alexander & Brown, 2010, 2011; Holroyd & Coles, 2002; Ullsperger et al., 2014). A prediction model is used to form an expectation about cues and outcomes (Niv & Schoenbaum, 2008; Ullsperger et al., 2014). A mismatch between the expected and observed result is used for updating the model to optimize performance-outcome predictions (Holroyd & Coles, 2002; Ullsperger et al., 2014). As such, conflicting external information is more likely to lead to internal model updating than concordant information (HajiHosseini et al., 2012; Ullsperger et al., 2014). These mismatches can be further exacerbated due to contextual effects, leading to misjudgments of advice accuracy (Hsee & Zhang, 2004; Ohmann et al., 2016). For instance, within the context of highly accurate information, moderately accurate advisors may be underestimated. Conversely, one’s expertise might be overestimated when other advisors turn out to be very inaccurate.

Information mismatches result in neurophysiological responses that can be observed with electroencephalography (EEG). First, an early event-related potential (ERP) component, the P2, is typically observed in fronto-central electrodes between 150 and 220 ms after stimulus presentation (Bellebaum et al., 20110131; Gheza et al., 2018; Holroyd et al., 2008; Martínez-Selva et al., 2019; Potts et al., 2006; San Martín et al., 2010; Williams et al., 2021; Wischnewski & Schutter, 2018, 2019). The P2 component is implicated in information and valence evaluation relative to the context (Glazer & Nusslock, 2022; Wischnewski & Schutter, 2018; Zou et al., 2022). As such, the P2 reflects an assessment of whether a cue or outcome is better or worse than expected compared to other available information (Potts et al., 2006; Wischnewski & Schutter, 2018, 2019). P2 is functionally correlated with a subsequent negativity, the N2, which is often identified as the feedback-related negativity (FRN; sometimes referred to as medial frontal negativity, MFN) (Hajcak et al., 2005, 2006, 2007; HajiHosseini et al., 2012; Holroyd & Coles, 2002; Kirsch et al., 2022; Peterburs et al., 2019; Williams et al., 2021; Wischnewski et al., 2018). The FRN is also observed in fronto-central electrodes and peaks between 200 and 350 ms after feedback onset. The dorsal anterior cingulate cortex and medial prefrontal cortex have been proposed as the cortical origin for FRN and P2 signals, which together may reflect activation of the mesocortical dopamine pathway (Alexander & Brown, 2011; Bayer & Glimcher, 2005; Hauser et al., 2014; Matsumoto et al., 2007; Meshi et al., 2012; Ridderinkhof et al., 2004; Ullsperger et al., 2014). Although this network is often related to reward processing (Haber & Knutson, 2010), the same brain regions hav been shown to encode social information (Behrens et al., 2008; Diaconescu et al., 2017; Konovalov et al., 2021). The FRN reflects processing of unexpected outcomes and considered the electrophysiological correlate of detecting prediction errors during performance monitoring (Cavanagh, Zambrano-Vazquez, et al., 2012; Cavanagh & Frank, 2014; Hajcak et al., 2006; Holroyd & Coles, 2002). Together, the P2-FRN complex is indicative of subjective valuation of decisions and outcomes. This may involve the evaluation of reward or punishing feedback or social information related to future feedback, including expert advice (Burnside & Ullsperger, 2020; Li et al., 2020; Wischnewski et al., 2018; Wischnewski & Schutter, 2019; Zou et al., 2022).

The second ERP component involved in performance monitoring and the evaluation of social information during decision making is the P3 (Polich, 2007; Ullsperger et al., 2014; Williams et al., 2021). The P3 is observed in fronto-parietal electrodes between 300-600 ms after stimulus presentation (Polich, 2007; Ullsperger et al., 2014). Depending on the task, it can occasionally be divided into an early fronto-central (P3a) and a later centro-parietal (P3b) component (Polich, 2007). The P3 is an attentional component and relates to conscious feedback processing (Novak & Foti, 2015; Ullsperger et al., 2014). Accordingly, it indicates an accumulation of evidence towards information being positive or negative. Consequently, the P3 is suggested to be involved in the maintenance and updating of internal prediction models (Huster et al., 2011; Walentowska et al., 2016). Further evidence suggests that P3 is involved in validation of social information (Li et al., 2020; Peterburs et al., 2019; Valt et al., 2020; Wischnewski et al., 2021; Wischnewski & Schutter, 2019; Yan et al., 2023). In line with this, the amplitude of the P3 component was larger when advice cues were accepted, compared to when they were rejected (Li et al., 2020). In other words, P3 may be an electrophysiological correlate of finetuning internal prediction models, when advice information is deemed to be informative.

Altogether, the role of P2, FRN and P3 in feedback processing has been thoroughly investigated and research has demonstrated that they play a significant role in social information processing (Boksem et al., 2011; Burnside & Ullsperger, 2020; Peterburs et al., 2019; Schutter et al., 2004; Valt et al., 2020; Zou et al., 2022), and specifically advice validity (Li et al., 2020; Wischnewski et al., 2018, 2021; Wischnewski & Schutter, 2019; Yan et al., 2023). Furthermore, amplitudes of these components are modulated when conflicting information is presented (Wischnewski & Schutter, 2019). However, to date it is unclear how these ERP signals relate to situations in which social information is interpreted erroneously. As alluded to above, depending on context advice validity can be either under– or overestimated. Such misjudgments are known as the distinction bias, which is the tendence to overly differentiate two or more options if they are presented together (Ceschi et al., 2019; Hsee & Zhang, 2004). For example, when judging the value of art, the advice of a student with an art major may be valued less in comparison to that of a renowned art expert. As such, the student’s advice may be (falsely) dismissed even if oneself is a layperson without prior knowledge. Conversely, the knowledge of a friend who visited an art gallery may be overestimated, when oneself and others have never done so. How performance monitoring signals like P2, FRN, and P3 are modulated by such misjudgments is currently unknown.

In two experiments the electrophysiological correlates of inaccurate processing of social information were investigated. To that end, we designed a decision task with multiple jointly presented advice cues, with high (*Experiment 1*) or low (*Experiment 2*) validity. In the first experiment all cues were similar in the sense that all had an accuracy above chance level, but distinct in that the level of accuracy differed. Analogously, in the second experiment all advise information was below chance level, but exact accuracies were dissimilar. Participants had the opportunity to learn about the advice information, with cues presented jointly. Subsequently, participants performed the same task with single cues to test whether they would follow or oppose the advisor. It was hypothesized that mismatches between cues would lead to generally increased amplitudes of performance monitoring signals. Furthermore, given the relationship between P3 and prediction model updating, we anticipated its amplitudes would be increased for more informative cues.

## Methods

### Participants

For *Experiment 1* thirty healthy adult were recruited (mean ± SD, 23.67 ± 3.36, 22 females). *Experiment 2* consisted of twenty-one healthy volunteers (mean ± SD, 22.57 ± 4.50, 16 females). For both experiments a power analysis was performed to estimate the sample size using G*power (Faul et al., 2009). A statistical power (β) of 0.95 and type I (α) error of 0.05 and correlation between session of r = 0.67 (Wischnewski et al., 2021). Studies which used the same protocol were not available before this study. Therefore, for *Experiment 1* an effect size of f = 0.25 (medium effect), based on previous experiments of lab using predictive social cues (Wischnewski et al., 2018, 2021; Wischnewski & Schutter, 2019). The calculation was based on a repeated-measures ANOVA and one independent variable with three conditions. This resulted in a suggested sample size estimate of N=30. For *Experiment 2*, sample size calculation was based on the approximate effect size in *Experiment 1* (f = 0.30), which resulted in a sample size estimate of N=21.

All participants were right-handed and had normal or corrected-to-normal vision. None of the participants used psychotropic medication or had a history of cognitive or neurological disorders. The study was approved by the local ethics committee of the Radboud University Nijmegen (The Netherlands).

### Decision making task

Participants were asked to indicate which of two vases is the most expensive. In previous studies, we demonstrated that participants experience this task as very difficult and without cues their performance is at chance level (Wischnewski et al., 2018, 2021; Wischnewski & Schutter, 2018, 2019). Vases were presented on a black screen (22 inch, 30×48 cm, resolution: 1680×1050) and participants were placed approximately 80 cm from the screen. The pictures of the vases (resolution: 350×250 pixels) had a monotonous light gray background and were presented 5 cm left and right from the center. Participants made their choice by pressing the left or right arrow key corresponding to the left or right vase. Each experiment consisted of three parts (**Fig. 1**). In the first part, the ‘initial phase’, participants were shown the vases for maximally 2000 ms and had to make a choice without any additional cues. Then, 500 ms after their decision, feedback was given by displaying the words ‘correct’ or ‘wrong’. If participants did not respond within the 2000 ms the word ‘faster’ was displayed. Feedback was shown for 1500 ms. The initial phase consisted of a total of 50 trials and lasted for approximately 3 minutes.

**Figure 1.**
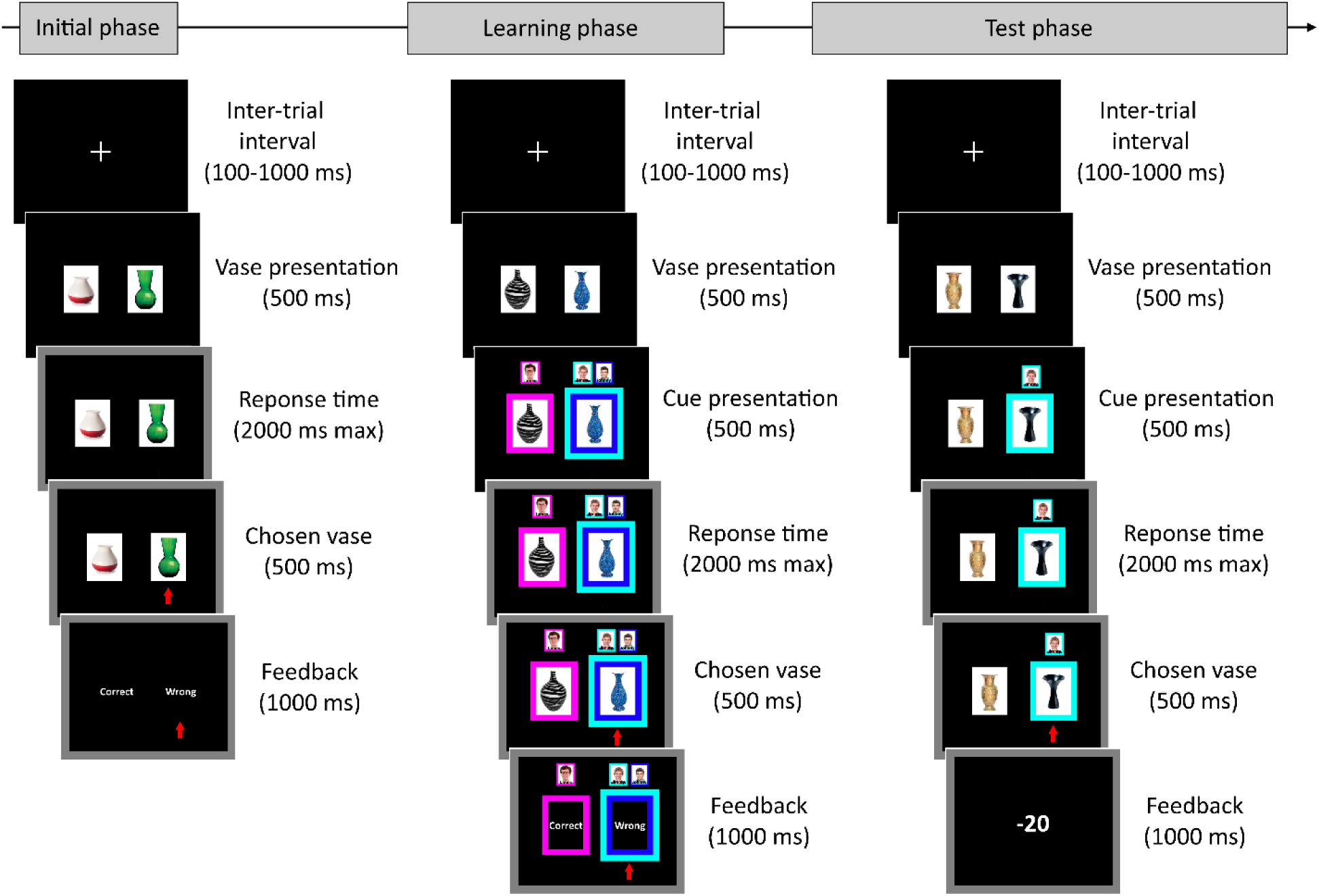
In both experiments an advice-cued decision making task was used that consisted of three stages. In all stages two vases were shown and participant had to indicate which of the two is most valuable. In the ‘initial phase’ participants did this without cues. In the ‘learning phase’ participants were presented with three advice cues after the vases were shown. In one condition all advisors agreed on the same vase and in the other three conditions one advisor cue deviated from the other two (as shown in the example in this figure). Cue validity differed per condition and per experiment. Cue validity in experiment 1 was 60%, 80% and 90%. In experiment 2 cue validity was 40%, 20%, and 10%. Finally, in the test phase participants played for points and were asked to score as high as possible. Unbeknownst to the participants, points and cues were presented pseudo-randomly. Only a single cue (of the previously learned ones) was presented to investigate the percentage following and decision.

The second part of each experiment was the ‘learning phase’. Vases were presented together with three simultaneous cues, which indicated the choices of three fictional players. Participants were instructed to learn how predictive each of the three cues is, as they would get to see these cues in the last phase in which they will play for points. Hence, the better they learned the predictive value, the better they could utilize these cues in the test phase. In *Experiment 1* the advice cues were highly predictive and indicated correct answers in 60%, 80% and 90% of the trials. This means that all cues were predictive for the correct answer above chance level (50%). In *Experiment 2* the advice cues indicated correct answers in 10%, 20% and 40% of the trials. This means that all cues had low predictive value, below chance level. In other words, that is the cues were more predictive for the wrong answer. Cues consisted of a colored (pink, cyan and cobalt blue) box around the vase, as well as a photo and name of the fictional players (‘Jimmy’, ‘Johnny’, ‘James’) above the box. Names and pictures of the advice cues were randomized before the experiment, but the predictive value of each cue was consistent throughout the learning phase. For example, if Johnny was corrected in 60%, Jimmy was correct in 80% and James was correct in 90% of the trials, this predictive value remained the same for the entire learning phase. With these accuracies, the three cues would select the same cue in 40% of trials. In approximately 34% of trials the 40/60% cues deviated from the other two cues. The 20/80% and the 10/90% cues deviated in 11% and 15% of trials respectively. Thus the 60% cue (*Experiment 1*) and 40% cue (*Experiment 2*) had the most deviating opinions. At the start of the learning phase participants were unaware of the predictive value of each cue. Vases together with cues were presented for maximally 2000 ms. Then, 500 ms after participants made their decision feedback followed with the words ‘correct’ or ‘wrong’ for 1500 ms. The learning phase consisted of 100 trials and lasted for approximately 7 minutes.

In the third part of each experiment, the ‘test phase’, vases were presented and in each trial one of the three previously learned cues was shown. The cues were displayed pseudo-randomly across trials. Participants were instructed that they could use their knowledge of the cues from the learning phase. In this phase of the task participants received points for their answers, and they were encouraged to score as many points as possible. The advice cue presentation and points system was used, validated and described in detail in previous studies (Wischnewski et al., 2018, 2021). Briefly, during the test phase advice cues were shown at random, meaning they had a predictive value of 50%. Thus, the cues predictive value did not match the learning phase. To prevent participants from realizing this ambiguous points system was used. Points ranged from –40 to +50 in steps of 10. Participants were informed that the gained points for choosing one vase are relative to the points they would have received if opted for the alternative. As the number of points for this alternative was not presented, participants were kept uncertain about the actual correctness of their choice. For example, participants chose the left vase and received 20 points. Yet, participants did not know if the other vase would have resulted in more or less than 20 points, meaning that they did not know if their answer was correct. This means that the points themselves were not indicative of their performance. This was done to prevent any learning or updating of internal prediction models during the test phase. Rather, to make their decisions participants had to rely solely on the information concerning the advice cues they gathered during the learning phase. Consequently, the amount of following each cue in the test phase is directly related to the biases towards or against an advisor formed during the learning phase. As in the other phases, vases together with the cue were depending on response time presented for a maximum of 2000 ms. Then, 500 ms after participants made their decision points feedback followed and was presented for 1500 ms. The test phase consisted of 150 trials (50 trials for each cue) and lasted for approximately 10 minutes.

*Experiment 1* and *Experiment 2* only differed on the predictability of the cues in the learning phase. The initial and test phase were identical in both experiments.

### ERP recording, preprocessing, and analysis

EEG was recorded during the task using an Active-Two system (BioSemi, Amsterdam, The Netherlands). Thirty-two electrodes were placed according to 10-20 system. EEG data was sampled at 2,048 Hz and a default online low pass filter (DC to 400 Hz) was applied. The data was stored for offline analysis using the Fieldtrip toolbox (Oostenveld et al., 2010) in Matlab (version 2020b, Mathworks, Natick, MA, USA).

EEG data was offline re-referenced to the average signal and was band-pass filtered between 1 and 30 Hz (48 dB/Octave). Ocular, muscular and other large artefacts were removed using independent component analysis (on average 2.47 and 2.62 components in *Experiment 1* and *2*, respectively). EEG segmentation was time-locked to the onset of the cues (cue-locked) and to the onset of the outcome (outcome-locked) during the learning phase (Burnside & Ullsperger, 2020). All epochs started 200 ms before the onset of the cue or feedback and ended 800 ms after the onset of the cue or feedback. The period from –200 to 0 ms was used for baseline correction and the data was demeaned. Next, data were visually inspected and remaining artefacts were removed (*Experiment 1*: 3.56%, *Experiment 2*: 2.47%).

As we used a revised version of our decision task (Wischnewski et al., 2018; Wischnewski & Schutter, 2019), we did not use a priori time bins for analysis. Instead, we used the cluster-based permutation model approach introduced by Maris and Oostenveld (Maris & Oostenveld, 2007). Monte-Carlo simulation with 1000 repetitions was used as the permutation method. The analysis was performed in the time window of 0 to 500 ms after cue or feedback presentation and the pooled signal of fronto-central electrodes (Fz, Fc1, Fc2, and Cz) was used (Wischnewski & Schutter, 2019).

It was hypothesized that behavioral effects would be driven by ERP in the learning phase, and we anticipated no signal differences in the test phase. No specific hypotheses were made for signals in the baseline phase. We used similar analysis steps as described above were used for these phases. ERP results of baseline phase and test phase are presented in **Supplementary Fig. 1-4**.

### Statistical Analysis

Behavior during the test phase of both experiments included the outcome measures: I) percentage following of cues, II) decision time, III) subjective predictive value (SPV). The latter reflects how informative a participant believes the cues to be. SPV is the absolute of the percentage following minus 50 (corresponding to chance level) normalized to 1 (Wischnewski et al., 2018). On this scale, cues that are either followed or opposed in all cases relate to a value of 1, as they are indicative of the to be taken action in each trial. However, cues that are followed at chance level provide no information and correspond to an SPV of 0.

Repeated measures analyses of variance (rmANOVA) were performed with percentage following and decision time as dependent variables. The independent variable was the cue validity (60%, 80%, and 90% in *Experiment 1*; 40%, 20%, and 10% in *Experiment 2*). For percentage following post-hoc analyses comprised of three one-sample t-tests (one for each cue) with the actual cue validity as test value. Additionally, paired samples t-tests were performed to compare percentage following between advice conditions. Similarly, for decision time post-hoc paired samples t-test were performed to compare the advice conditions.

Cluster-based permutation analyses were performed to examine differences in cue-locked and outcome-locked ERP signals in the learning phase. Four conditions were compared: I) all cues point towards the same vase, II) 80% and 90% cue indicate one vase and the 60% cue points towards the other vase (60% deviant), III) 60% and 90% cue indicate one vase and the 80% cue points towards the other vase (80% deviant), IV) 60% and 80% cue indicate one vase and the 90% cue points towards the other vase (90% deviant). To confirm cluster statistics, significant results were followed by a rmANOVA with the average ERP amplitude within the cluster time window as dependent variable, and the four conditions as independent variable. This was followed up by post-hoc paired t-tests, comparing the four conditions.

In addition to the frequentist analyses, Bayesian one-sample and paired t-tests were performed to augment post-hoc results (Rouder et al., 2009). Default priors following a Cauchy distribution, with the center at 0 and a width of 0.707 were used (Vehtari et al., 2021). We report the Bayes factor (BF) that indicates the probability of the alternative hypothesis being more likely than the null hypothesis (BF_10_). BF_10_ > 3 is typically viewed as an indication that evidence points towards the alternative hypothesis, whereas BF_10_ < 0.33 points towards the null hypothesis (Rouder et al., 2009).

Finally, Pearson correlations were performed on the data of both experiments combined. We correlated the SPV (regardless of cue validity) with decision time, as well as P2 and P3 amplitude. For the latter, ERP amplitudes of deviating conditions were baseline corrected by subtracting the amplitudes of the control condition (no deviating cues).

All statistical analyses were performed using JASP (version 0.14.0.0) and custom scripts in Matlab. Throughout the manuscript mean values are shown with the standard error of mean (SEM).

## Results

### Experiment 1: Behavioral responses to highly accurate cues

During the learning phase participants were presented with cues that were correct in 60%, 80% and 90% of trials. During the test phase participants could use these cues to guide their decision. Since rewards were uninformative during the test phase, we anticipated that any biases towards or against cues would form during the learning phase. Furthermore, it was hypothesized that due to a framing effect the cue that seems most different from the others would be misjudged. Since the 60% cue showed the most deviating opinions compared to the highly valid 80% and 90% cue (see methods), we expected that the actual predictive value of the 60% cue would be underestimated. In addition, we also investigated the subjective predictive value (SPV), reflecting how informative participants judged a cue to be. A high SPV reflects a cue that is highly informative for the participant in making their decision (agree or oppose). A low SPV reflects a cue that does not relate clearly to a subsequent action.

As hypothesized, the percentage following differed significantly for the three cues (F(2,58) = 107.19, p < 0.001; **Fig. 2A**). Results showed that the 90% cue was followed in 93.12 ± 1.19% of trials, which was slightly more often than is expected from the objective predictive value (t(29) = 2.62, p = 0.014, d = 0.478, BF_10_ = 3.39). The 80% cue was followed in 76.10 ± 3.33% of trials, which was not significantly different than would be expected from the objective predictive value (t(29) = 1.17, p = 0.251, d = 0.214, BF_10_ = 0.36). The 60% cue was followed in 31.51 ± 3.48% of trials, which was significantly less than would be expected from the objective predictive value (t(29) = 8.18, p < 0.001, d = 1.493, BF_10_ = 2.44e^6^). Additionally, we found that decision time in the test phase was significantly affected by cue validity (F(2,58) = 23.13, p < 0.001; **Fig. 2B**). Interestingly, decision time followed the same pattern as SPV (**Fig. 2C**). Participants reacted fastest after the cue which they believed was most informative (90% cue), and slowest for the cue which they believed to be the least informative (60% cue).

**Figure 2.**
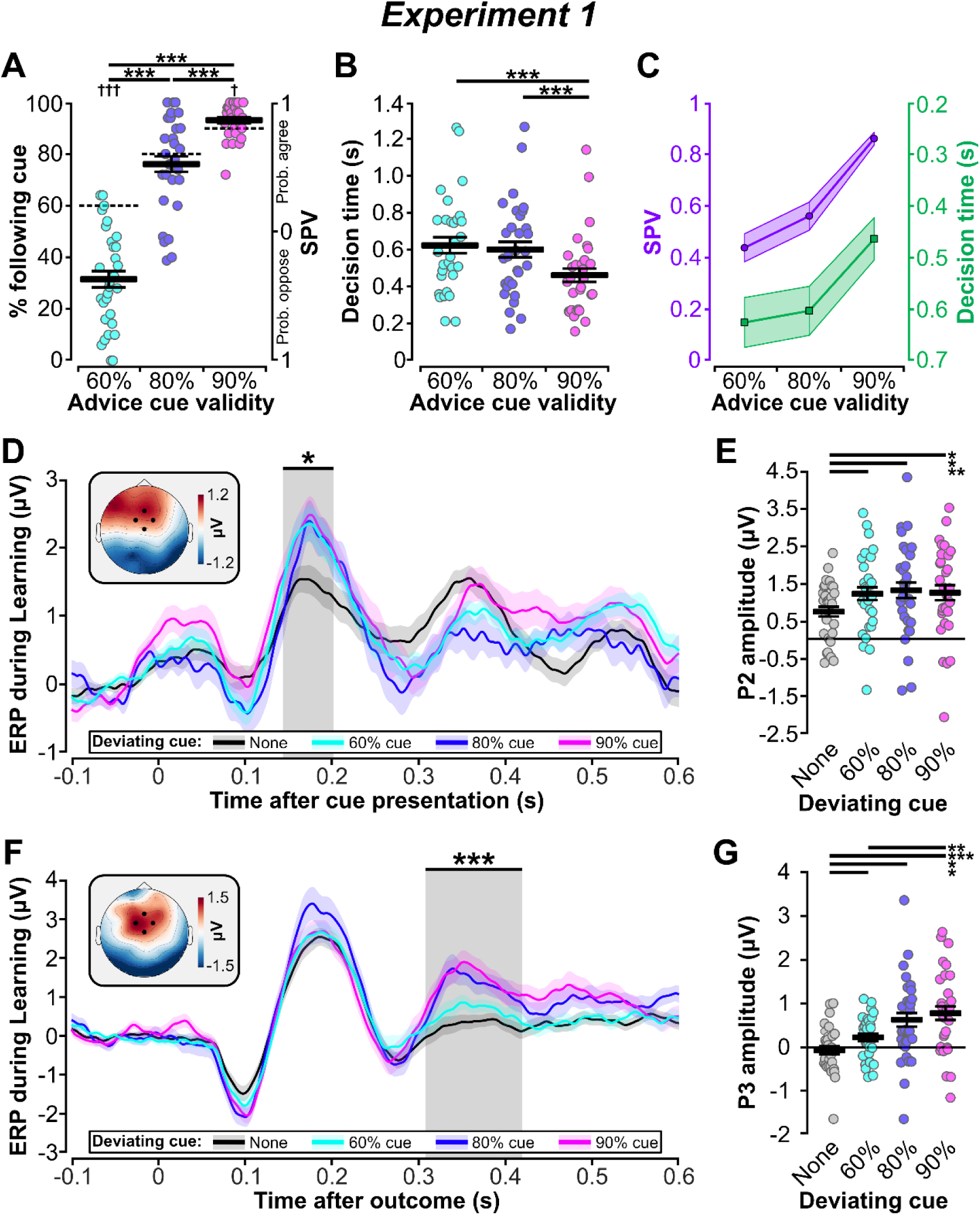
Behavioral (test-phase) and ERP (learning phase) results for Experiment 1. A) Individual and average percentage following and SPV, and B) Decision time. * Represents significant pairwise differences, † represents significant one sample t-test results. C) A comparison of general patterns of SPV and decision time. D) Cue-locked ERP signals for the four advice cue conditions. The topographical plot represents the P2 difference between deviant cues and all cues together conditions (time window: 136-202 ms). E) Individual and average P2 amplitudes per condition. F) Outcome-locked ERP signals for the four advice cue conditions. The topographical plot represents the P3 difference between the 90% cue condition (pink) and all cues together condition (black; time window: 309-424 ms). G) Individual and average P3 amplitudes per conditions.

### Experiment 1: Electrophysiological responses to highly accurate cues

During the learning phase participants had the opportunity to study the three cues and judge their validity, which could be used to guide choices during the subsequent test phase. Consequently, we expected that differences in cue-related signals during the learning phase may be indicative of behaviors observed in the test phase. During the learning phase we looked at two time points: I) when the cues were presented (cue-locked), and II) when feedback on the accuracy of the cues was provided (outcome-locked).

First, a cluster-based permutation test was performed on signals relating to four conditions (cues all together, 60% cue deviates, 80% cue deviates, 90% cue deviates) during cue presentation. A positive cluster (clusterstat = 1961.13, p = 0.019) was identified for the time window of 136-202 ms after stimulus presentation. This deflection corresponds to frontocentral P2 (**Fig. 2D**). This observation was confirmed by a separate rmANOVA on the signals within the same time window F(3,87) = 4.18, p = 0.008. Post hoc t-tests (**Fig. 2E**) showed significant differences between the cues together condition and the 60% cue deviant (t(29) = 3.36, p = 0.002, d = 0.613, BF_10_ = 16.56), 80% cue deviant (t(29) = 2.78, p = 0.010, d = 0.507, BF_10_ = 4.69) and 90% cue deviant

(t(29) = 2.85, p = 0.008, d = 0.521, BF_10_ = 5.48) conditions.

Second, a cluster-based analysis was done on outcome-locked signals. A positive cluster (clusterstat = 4030.57, p = 0.002) was found for the time window of 309-424 ms after feedback presentation, corresponding to the P3 component (**Fig. 2F**). We confirmed this with rmANOVA on this time window (F(3,87) = 9.11, p < 0.001). Post hoc t-tests (**Fig. 2G**) showed significant differences between the cues together condition and the 60% cue deviant (t(29) = 3.07, p = 0.005, d = 0.560, BF_10_ = 8.62), 80% cue deviant (t(29) = 3.03, p = 0.005, d = 0.554, BF_10_ = 8.05) and 90% cue deviant (t(29) = 4.53, p < 0.001, d = 0.828, BF_10_ = 278.62) conditions. Additionally, a significant difference between the 60% and 90% cue deviant condition was found (t(29) = 3.45, p = 0.002, d = 0.630, BF_10_ = 20.32). In summary, P3 was modulated by the validity of deviating cues. As the P3 component is related to performance monitoring, it could suggest that objective accuracy of unique cue information relates to the updating of one’s predictive model.

### Experiment 2: Behavioral responses to low accurate cues

In *Experiment 2* the effect of low predictive cues was investigated. As in *Experiment 1*, we hypothesized that the most deviating cue (40% cue in this case) would result in misjudgment of its accuracy. We found that participants followed all cues significantly more than is expected from the objective predictive value, with the largest effect for the 40% advice cue (F(2,40) = 34.14, p < 0.001; **Fig. 3A**). Specifically, the 10% cue was followed 26.19 ± 5.08% of trials (t(20) = 3.19, p = 0.005, d = 0.695, BF_10_ = 9.61). The 20% cue was followed in 45.96 ± 6.61% of the trials (t(20) = 3.93, p < 0.001, d = 0.857, BF_10_ = 42.32). The 40% cue was followed in 85.96 ± 4.31% of the trials (t(20) = 10.66, p < 0.001, d = 2.33, BF_10_ = 1.02e^7^). Decision time in the test phase was significantly affected by cue validity (F(2,40) = 6.02, p = 0.005; **Fig. 3B**). As in *experiment 1*, decision time followed a similar pattern as SPV (**Fig. 3C**).

**Figure 3.**
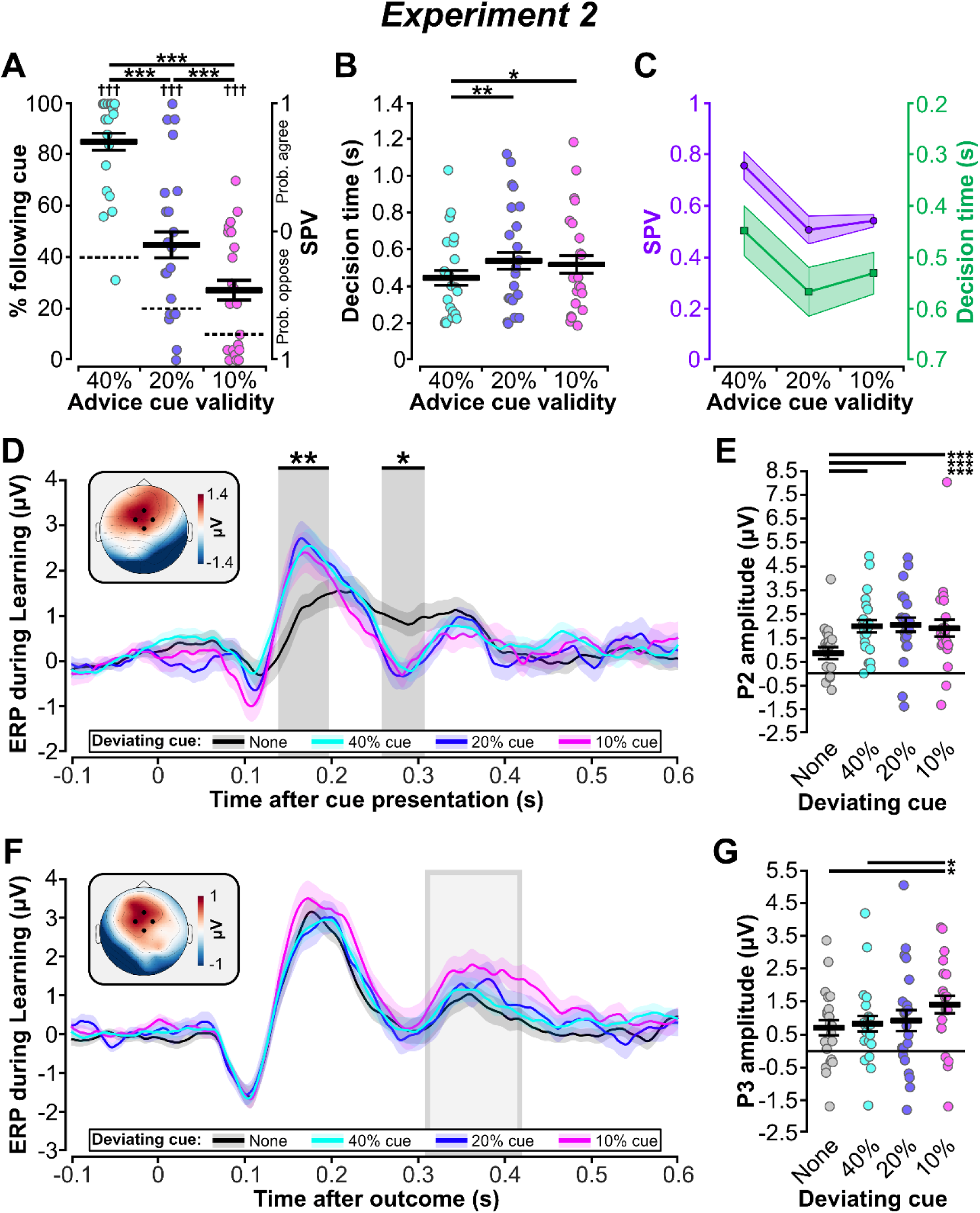
Behavioral (test-phase) and ERP (learning phase) results for Experiment 2. A) Individual and average percentage following and SPV, and B) Decision time. * Represents significant pairwise differences, † represents significant one sample t-test results. C) A comparison of general patterns of SPV and decision time. D) Cue-locked ERP signals for the four advice cue conditions. The topographical plot represents the P2 difference between deviant cues and all cues together conditions (time window: 131-194 ms). E) Individual and average P2 amplitudes per condition. F) Outcome-locked ERP signals for the four advice cue conditions. The topographical plot represents the P3 difference between the 10% cue condition (pink) and all cues together condition (black; time window: 309-424 ms). G) Individual and average P3 amplitudes per conditions.

### Experiment 2: Electrophysiological responses to low accurate cues

As in *Experiment 1* we inspected the ERP signals during the learning phase, after cue presentation (cue-locked) and after receiving feedback (outcome-locked). First, a cluster-based permutation test was performed on signals relating to four conditions (cues all together, 40% cue deviates, 20% cue deviates, 10% cue deviates) during cue presentation. Two clusters cluster were found between 131-194 ms (clusterstat = 3280.28, p = 0.006) and 260-310 ms (clusterstat = 1847.43, p = 0.028) after stimulus presentation respectively (**Fig. 3D**). These deflections correspond to the P2 and subsequent N2 (or FRN) component. As in *Experiment 1* the P2 for the deviant cues was significantly larger compared to the condition where all cues are presented together (rmANOVA on the P2 time window: F(3,60) = 7.65, p < 0.001). Post hoc t-tests (**Fig. 3E**) showed significant differences between the cues together condition and the 40% cue deviant (t(20) = 5.06, p < 0.001, d = 1.10, BF_10_ = 437.43), 20% cue deviant (t(20) = 3.91, p < 0.001, d = 0.854, BF_10_ = 41.21) and 10% cue deviant (t(20) = 4.25, p < 0.001, d = 0.927, BF_10_ = 81.57) conditions.

A second cluster-based analysis on outcome-locked ERP signals did not reveal any significant clusters. However, we performed an rmANOVA comparing the four conditions within the P3 time window from *Experiment 1* (309-424 ms). The results suggested a significant difference between P3 signals (F(3,60) = 3.441, p = 0.022; **Fig. 3F**). Post-hoc t-tests (**Fig. 3G**) showed a significant difference between the cues together condition and the 10% cue deviant conditions (t(20) = 2.66, p = 0.015, d = 0.582, BF_10_ = 3.64), as well as between the 40% cue deviant and 10% cue deviant conditions (t(20) = 2.67, p = 0.015, d = 0.583, BF_10_ = 3.66).

### Correlation analysis on the SPV

For a correlation analysis we combined the data of all conditions in both experiments. First, we found a significant negative correlation between SPV and decision time (r = 0.374, p < 0.001; **Fig 4A**). This indicates that a cue that was perceived as more informative also related to faster decision. Next, we correlated SPV with cue-locked P2 (**Fig. 4B**) and outcome-locked P3 amplitudes (**Fig. 4C**) for when one cues deviates from the other, corrected for when all cues are shown together. This correlation indicates whether a mismatch during cue presentation is related to larger ERP amplitudes when the advice cue is viewed as informative. Indeed, small but significant correlations were found for P2 (r = 0.184, p = 0.024) and P3 (r = 0.171, p = 0.034), suggesting that amplitudes of both components are indicative of subjectively perceived advice validity.

**Figure 4.**
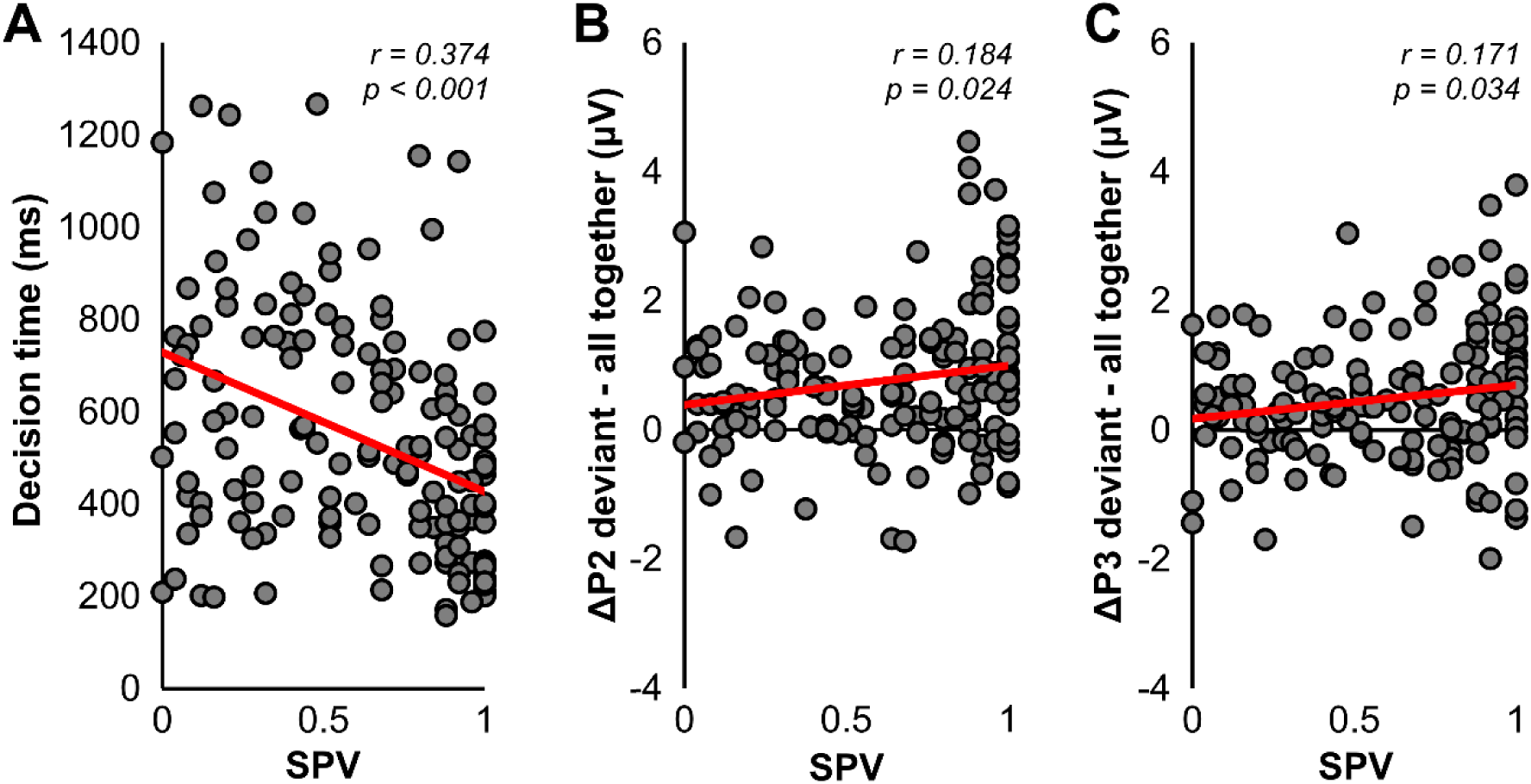
Scatterplots showing the correlation between SPV and A) decision time, B) P2 amplitude of disagreeing advisors controlled for the condition in which all advisors agree, and C) P3 amplitude of disagreeing advisors controlled for agreeing advisors.

## Discussion

The aim of this study was to identify the electrophysiological correlates judging and misjudging validity of social information. In two experiments participants compared different advice cues and judged their validity in a decision making task. Comparison of social information of similar but differing accuracy can lead to a distinction bias, which results in a systematic under– or overestimating of advice validity. During cue presentation the amplitude of the P2 component was increased when one advisor deviated from the others. Furthermore, we found that the P3 component during feedback presentation, that indicated the correctness of the cues, was modulated by different advice cue conditions. Specifically, when all advisors agreed the P3 amplitude was smallest. In the case that one advisor’s opinion did not match with the others, a linear increase in P3 was observed with the largest amplitudes for the most informative advice cue. In agreement with this observation, we found a significant positive correlation between SPV and P2 as well was P3, regardless of which advice cue was presented. Together, these data suggest that P2 and P3 components are elevated during a social information mismatch and are modulated by the validity of the information.

When inspecting the behavioral responses, we observed that within the context of highly accurate advice, a cue with moderately high accuracy was underestimated. Participants opposed this cue, even though the optimal strategy would have been to follow the cue. Conversely, within a context of advice with low accuracy, participant overestimated a moderately inaccurate cue. They followed the cue although the optimal strategy would be to oppose it. These observations are in line with the so-called distinction bias (Ceschi et al., 2019; Hsee & Zhang, 2004). When (social) information is presented together we have the tendency to overemphasize existing differences. In *Experiment 1* we used highly accurate cues with validities of 60%, 80% and 90%. So, the correctness of each cue is above chance level (50%). As participants experienced the judging of value of vases nearly impossible, they scored at chance level. Consequently, the optimal strategy would be to follow all cues. However, due to the distinction bias the 60% cue was perceived as inaccurate compared to the others and was thus believed to be less valid than it really was. It was only followed in ∼32% of trials. In *Experiment 2* all advice validities were below chance level (40%, 20% and 10%), meaning that the optimal strategy is to oppose these cues. Again, due to the distinction bias the 40% cue was perceived as better than it factually was and was followed in ∼85% of the trials.

Concerning the electrophysiological data, we found that when participants observe disagreement among advisors, increased P2 amplitudes were observed. This result was consistent across conditions and both experiments. This suggests that the P2 is involved in processing conflicting social information, and extends findings in previous studies which indicated that the P2 is involved in conflict monitoring (Holroyd et al., 2008; Wischnewski & Schutter, 2018; Xiao et al., 2019; Zinchenko et al., 2015, 2017). Further, in a previous study we found that the P2 amplitude was modulated by the advice of an expert (Wischnewski & Schutter, 2019). Specifically, an interaction between future reward or punishment and expert agreement or disagreement with one’s own opinion was found. Although the task used in our previous study was considerably different from the task used here, it does show that early fronto-central activity is crucially involved in the assessment of social information. Moreover, P2 amplitude was positively correlated with how informative a cue was perceived independent of its actual accuracy. For this we calculated the SPV (Wischnewski et al., 2018), which is a metric where a value of 1 indicates that a cue is subjectively highly informative. This can either mean that participants always follow or oppose the advice. A value of 0 on the other hand indicates that the cue is followed at chance level and is therefore uninformative. We found a small but significant positive correlation between P2 and SPV. This suggests that the P2 is related to subjective levels of certainty that advice is accurate or not. In line with this, Gole et al. (Gole et al., 2012) found that P2 during an affective cueing paradigm is modulated by certainty and that individuals with low tolerance for uncertainty had larger P2 amplitudes. Generally, evidence from this study and previous studies indicates that the P2 is an early component in the performance monitoring process which highlights mismatches between cues. Although the P2 may not be directly involved in internal model updating, it may initiate the process by flagging cues with potentially significant informational value.

In *Experiment 2*, but not in *Experiment 1*, we observed a modulation of the FRN following the P2 component. The effect for FRN was the same as for P2, with a significantly larger negativity for disagreeing advise cues as compared to advice cues in agreement. This is in line with the observation that the FRN (or N2) is involved in mismatch detection and processing of unexpected information (Cavanagh et al., 2010; Cavanagh, Zambrano-Vazquez, et al., 2012; Cavanagh & Frank, 2014; Hajcak et al., 2005, 2006; HajiHosseini et al., 2012; Holroyd & Coles, 2002). Although P2 and FRN are functionally distinct, they are correlated and reflect similar and overlapping processes (Cavanagh, Figueroa, et al., 2012; Holroyd et al., 2008). Previously, larger FRNs have been demonstrated in relation to outcomes that were unexpected based on the provided social information (Wang et al., 2020; Wischnewski & Schutter, 2019). However, with the absence of consistent modulation of the FRN in this study, the role of this ERP component in social information misjudgment remains unclear and requires further investigation.

Similar to P2, we observed that disagreement among advisors resulted in increased P3 amplitudes. However, in contrast to P2, P3 was differentially modulated depending on which advice cue deviated from another. This modulation was linearly related to the objective informative value of the cue. Specifically, in *Experiment 1* the 90% cue, which is highly informative as it is almost always correct – so, it should be followed – resulted in the largest P3, whereas the 60% cue, which is objectively the most uncertain resulted in the lowest P3 amplitudes. Analogously, in *Experiment 2*, the 10% cue, which is also highly informative as it almost always wrong – so, it should be opposed – resulted in the largest P3 amplitude, whereas the objectively most uncertain 40% cue resulted in the smallest P3 amplitudes. Together this suggests that the P3 component does not directly reflect accuracy, but rather how much information can be extracted from a deviating advice cue. A similar observation was made when considering the subjective valuation of advice accuracy. P3 was significantly positively correlated with SPV, meaning that when participants rated a cue as informative, this was reflected by increased P3 amplitude. Together these findings seem to be in agreement with the idea that P3 reflects updating of internal prediction models.

Furthermore, the observation that uncertain advice information (40% and 60% cues) was related to the smallest P3, may explain why the accuracy of these cues was most likely to be misjudged. If P3 amplitude is positively correlated to belief updating (Huster et al., 2011; Jepma et al., 2018; Ullsperger et al., 2014; Walentowska et al., 2016), small amplitudes may be associated with erroneous or equivocal prediction model revision (Li et al., 2020). Our observation is in agreement with the that of Li et al (Li et al., 2020), who showed decreased P3 amplitudes following incorrect advice and when advice was rejected. Together, this suggests that P3 relates to two processes: I) Judgement of social information accuracy, and II) behavioral adaptation to follow or disregard the advice. Concerning the misjudgment of advice accuracy, we propose that the distinction bias is related to smaller P3 amplitudes, which are associated with suboptimal performance monitoring.

P3 can be divided into two sub-components, namely the P3a, which is most dominant in fronto-central electrodes, and P3b, which is most dominant in centro-parietal electrodes (Fischer & Ullsperger, 2013; Polich, 2007). Topographic plots (Figure 2F, 3F) suggested that advice-related differences in ERP signals were mainly observed in fronto-central areas, suggesting modulation of P3a. However, due to the absence of a clear deflection in parietal electrodes we were unable to distinguish both sub-components of P3. Both P3a and P3b are involved in prediction model updating. Whereas P3a is an attention-related component that is associated with conscious processing of feedback data (Polich, 2007; Ullsperger et al., 2014), P3b is related to behavioral adaptions in response to undesirable outcomes (Chase et al., 2011; Fischer & Ullsperger, 2013; Grahek et al., 2023; Ullsperger et al., 2014). Although we demonstrated modulation of the P3 amplitude, the current data do not allow us to make inferences on whether (mis)judgment mostly relates to P3a, P3b, or both.

One notable observation is that in *Experiment 2* with the low accuracy cues, cue validity was generally overestimated. The opposite behavioral pattern, that is all cues being underestimated, was however not observed in *Experiment 1* with the high accuracy cues. This suggests high levels of uncertainty during decision making, to the point that even very low-accurate advisors are trusted. As such, misjudging social information in this study cannot solely be explained by the distinction bias, but may also include others, like the bandwagon effect and the trust bias. The bandwagon effect is the tendency to conform to the opinions of others (Bindra et al., 2022), whereas the trust bias describes the preference for believing that statements of others are truthful (Street & Masip, 2015). The present study was not designed to highlight such biases. Therefore, we cannot draw conclusions about the involvement of advice-related ERP signals in these and other biases and may be an interesting topic to address in future studies.

## Conclusions

Social information can help us to make decisions in uncertain situations. However, our ability to judge whether such social information is accurate relies on heuristics and can lead to cognitive biases. The consequence can be a significant misjudgment of the validity of social cues. In this study we show that processing of advice cues relates to the modulation of the P2 component, when the cues are presented, and the P3, when subsequent outcomes are presented. P2 reflects a mechanism for detecting agreement or disagreement between advisors, with increased amplitude for the latter. The P3 is coupled to a process of assessing accuracy of advice and belief updating. We propose that small P3 amplitudes reflect erroneous assessment of advice cues, which can yield misjudgments and biased interpretation of social information.

