## Supplementary Figure for "Electrophysiological correlates of (mis)judging social information"

**Supplementary Materials**

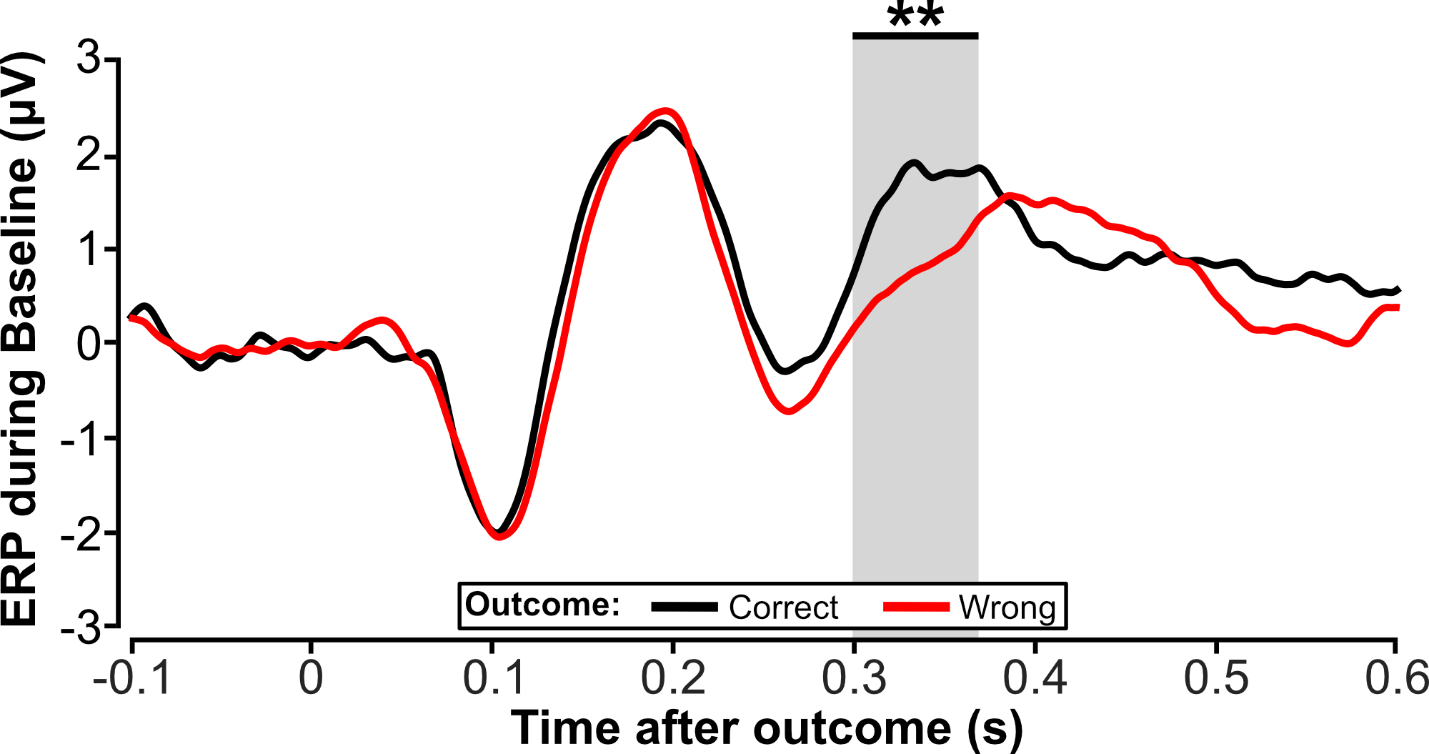

Supplementary Fig 1. ERPs after feedback presentation during baseline phase of *Experiment 1* (no cues presented). Feedback indicating correct responses is represented by the black line, whereas feedback indicating wrong responses is represented by the red line. A difference was observed between ERP signals related to positive and negative outcomes in the time between 297 and 369 ms after feedback presentation (clusterstat = 1436.30, p = 0.008).

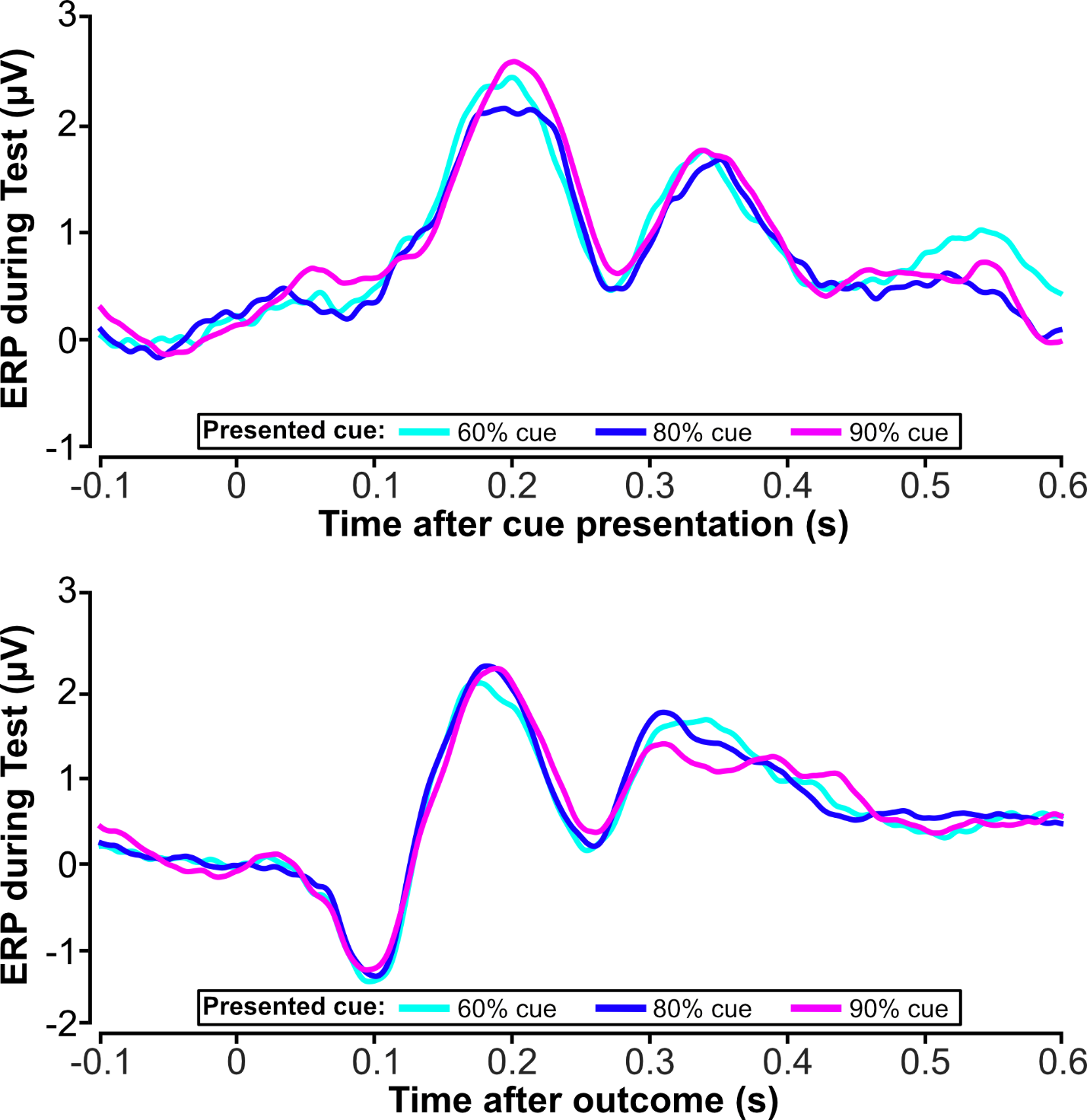

Supplementary Fig 2. ERPs after cue (upper panel) and outcome (lower panel) presentation during test phase of *Experiment 1*. In each trial one cue was presented and ERP responses for each cue are shown. No significant differences were observed.

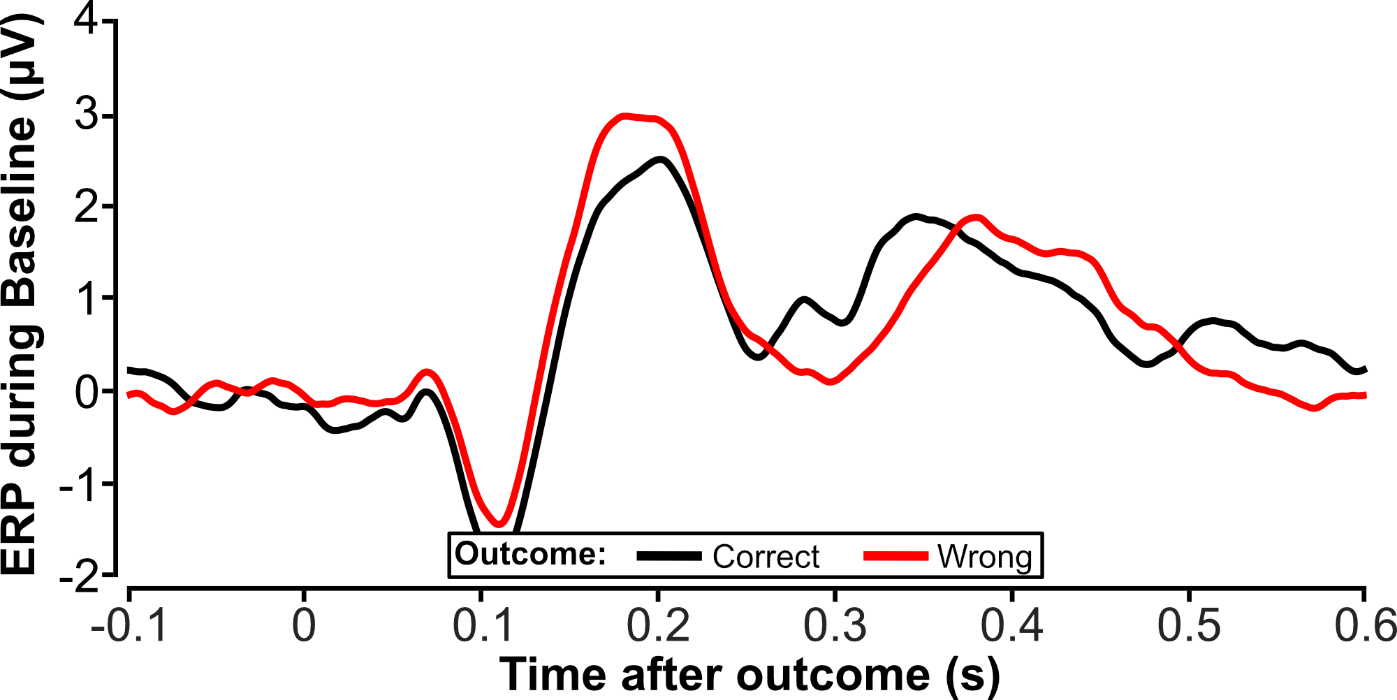

Supplementary Fig 3. ERPs after feedback presentation during baseline phase of *Experiment 2* (no cues presented). Feedback indicating correct responses is represented by the black line, whereas feedback indicating wrong responses is represented by the red line. No significant differences were observed.

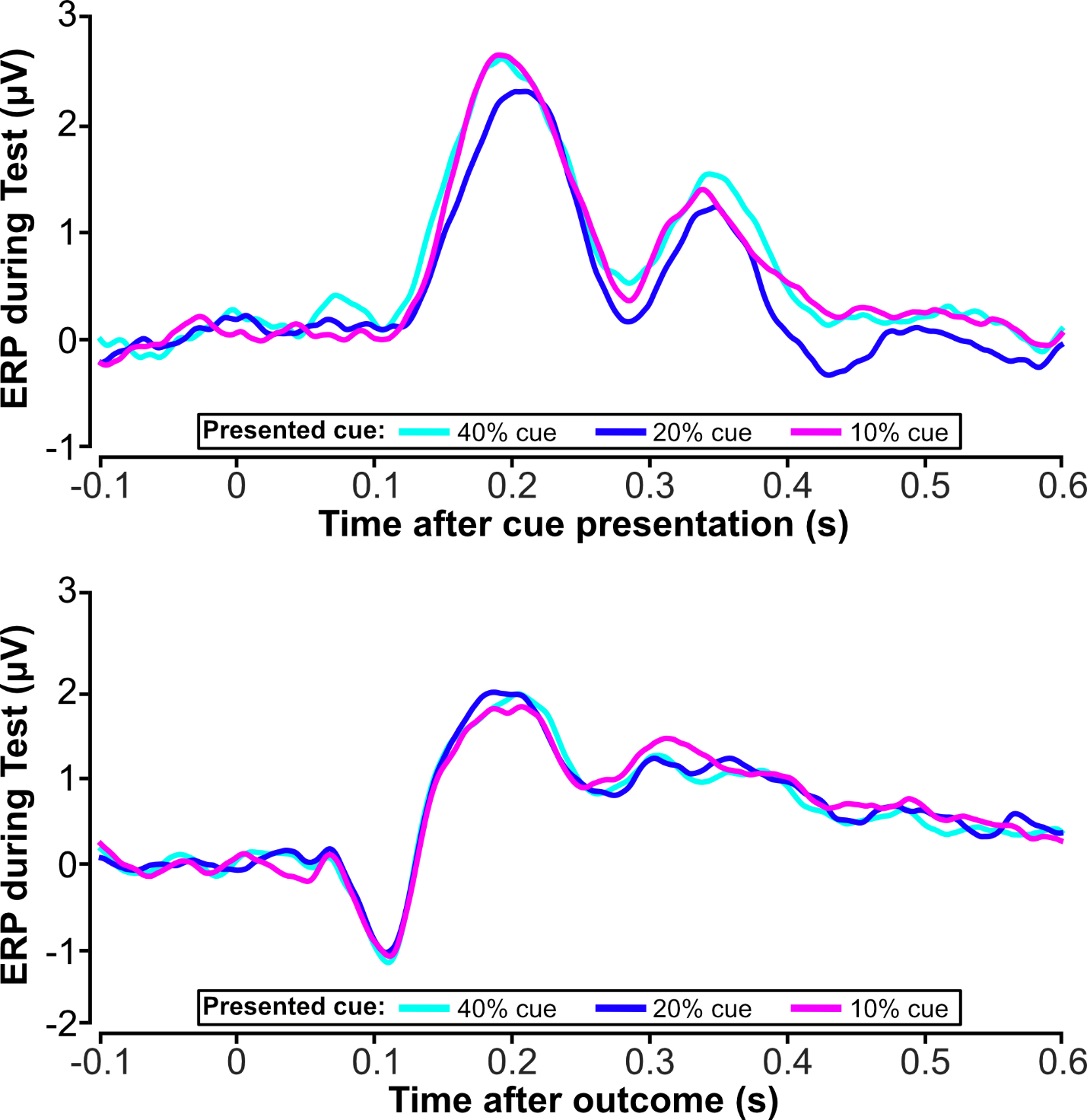

Supplementary Fig 4. ERPs after cue (upper panel) and outcome (lower panel) presentation during test phase of *Experiment 2*. In each trial one cue was presented and ERP responses for each cue are shown. No significant differences were observed.
